# Stage-specific exposure to an activity-permissive media enhances neuronal maturation in oligodendrocyte-enriched cortical organoids

**DOI:** 10.64898/2026.05.17.725797

**Authors:** Clara Chung, Michelle J. Kim, Grace Field, Konstantina Pilarinos, Elizabeth K. Kharitonova, Natalie Baker Campbell, Christopher V. Gabel, Joseph L. Orofino, Ella Zeldich

**Affiliations:** Department of Anatomy & Neurobiology, Boston University Chobanian & Avedisian School of Medicine, Boston University, Boston, MA; Department of Biological Sciences, CUNY Hunter College, New York, NY 10065; Department of Physiology and Biophysics, Boston University Chobanian & Avedisian School of Medicine, Boston University, Boston, MA; Neurophotonics Center, Boston University, Boston, MA; Bioinformatics Program, Boston University, Boston, MA, 02118; Department of Medicine, Boston University Chobanian & Avedisian School of Medicine, Boston University, Boston, MA

**Keywords:** organoids, oligodendrocytes, oligodendrocyte-enriched cortical organoids, neuronal activity, BrainPhys, calcium imaging, myelination models, neuron-glia interactions, cortical development

## Abstract

Oligodendrocyte-enriched cortical organoids (OCOs) are a powerful platform for modeling oligodendrogenesis in a human cellular context. However, neuronal activity is impaired in conventional culture media, limiting assessment of neuronal function in conjunction with oligodendrocyte biology. To address this, we used a modified BrainPhys medium termed neuronal activity medium (NAM) and defined the optimal developmental window for NAM exposure to generate OCOs with robust neuronal activity (NAM-OCOs). Stage-specific exposure to NAM, prior to oligodendrocyte expansion, leads to enhanced structural maturation, as evidenced by increased organoid size, heightened synaptogenesis, and upregulation of transcripts associated with neuronal complexity. Further, NAM-OCOs display increased cellular heterogeneity, including greater representation of GABAergic interneurons while preserving oligodendrocyte development and maturation. Altogether, our studies demonstrate that stage-specific exposure to an activity-permissive environment enhances neuronal activity, establishing an OCO model which integrates neuronal activity with oligodendrocyte development and maturation.

**Highlights:**

- Increased neuronal activity in oligodendrocyte-enriched cortical organoids (OCOs)
- Stage-specific Neuronal Activity Medium (NAM) optimizes activity
- NAM-OCOs display increased cellular heterogeneity and neuronal maturation
- Oligodendrogenesis is preserved in NAM-OCOs

**eTOC blurb:** In this article, Chung et al enhance neuronal activity in oligodendrocyte-enriched cortical organoids (OCOs) through stage-specific exposure to Neuronal Activity Medium (NAM). OCOs exposed to NAM display elevated cellular heterogeneity, structural maturation, and synaptogenesis, while preserving oligodendrocyte development and maturation. These results establish an increasingly comprehensive OCO model for studying neuronal function and oligodendrogenesis.

## Introduction

Oligodendrocytes are the myelinating cells of the central nervous system (CNS), providing axonal insulation and metabolic support to neurons (Fünfschilling et al., 2012). Much of our foundational understanding of oligodendrocytes has been informed by rodent models (Bradl and Lassmann, 2010). However, knowledge of the developmental and maturation trajectories of oligodendrocytes in a human genetic context remains limited and largely inferred from postmortem tissue. While numerous methods have been established for generating oligodendrocytes from human induced pluripotent stem cells (iPSCs) (Douvaras et al., 2014; García-León et al., 2018; Wang et al., 2013), as with other neural cell types, a 2D environment fails to capture the diverse cellular heterogeneity, microenvironment, and complex oligodendrocyte-neuron interactions present in the *in vivo* milieu (Zeldich and Rajkumar, 2024).

The rise of iPSC-derived brain organoids has transformed modeling of human brain development. By harnessing the intrinsic self-organizing capabilities of iPSCs, these three-dimensional (3D) cellular models capture previously inaccessible stages of brain development. Within organoids, diverse cell types, including neural progenitors, astrocytes, and diverse neuronal subtypes co-develop in a coordinated manner, recapitulating complex cellular interactions (Kadoshima et al., 2013; Lancaster et al., 2013; Paşca et al., 2015; Velasco et al., 2019). Beyond structural features, organoids display spontaneous synaptic activities, assemble functional networks, and recapitulate electrophysiological properties characteristic of mid-gestational neurons (Fitzgerald et al., 2024; Paşca et al., 2015; Sharf et al., 2022; Trujillo et al., 2019; Yakoub, 2019). The feasibility of functional analyses in organoids thus provides an excellent platform for modeling early-stage electrical activity and network dynamics in a human context.

Human neurodevelopment is dictated by the complex interplay between multiple, functionally distinct cell types. While early organoid models excluded oligodendrocytes, several approaches have emerged to promote the development of oligodendrocytes in organoids (Madhavan et al., 2018; Marton et al., 2019; Shaker et al., 2021). In particular, work by (Madhavan et al., 2018) demonstrated exposure to platelet-derived growth factor AA (PDGF-AA) and insulin-like growth factor 1 (IGF-1) followed by thyroid hormone (T3) after neuronal expansion supports the formation of oligodendrocyte progenitor cells (OPCs) and mature, myelinating oligodendrocytes. This model omits ventralizing or caudalizing morphogens, including sonic hedgehog (SHH) and retinoic acid (Qi et al., 2002), producing oligodendrocyte-enriched cortical organoids (OCOs) with a dorsal telencephalic identity. These OCOs recapitulate the distinct developmental origin of oligodendrocytes derived from dorsal cortical progenitors and enable investigation of oligodendrocyte maturation within a cortical neuronal environment.

While OCOs provide an excellent platform for modeling oligodendrocyte development, the protocol established in (Madhavan et al., 2018) describes prolonged culture in a conventional basal medium. While traditional media such as Neurobasal medium and DMEM support the long-term growth and survival of neuronal cultures, consistent with our findings, they do not support spontaneous neuronal activity (Bardy et al., 2015). As a result, functional modeling of neuron-oligodendrocyte interactions in a dorsal organoid system is limited.

BrainPhys is a chemically defined neuronal medium developed to bridge the gap between traditional culture conditions and the extracelllular environment *in vivo* and provides an increasingly physiologically relevant environment supportive of spontaneous neuronal activity and long-term growth. Furthermore, by promoting spontaneous activity, BrainPhys indirectly enhances neuronal maturation (Bardy et al., 2015; Satir et al., 2020).

Given the importance of electrical activity in neuronal and oligodendrocyte development, we investigated the effects of long-term maintenance of OCOs in an environment permissive to spontaneous activity. Using a modified BrainPhys formulation, termed here as Neuronal Activity Media (NAM), we defined the optimal developmental time window, during which exposure to an activity-permissive environment maximally enhances neuronal activity in OCOs. We then profiled the cellular and molecular features of OCOs with enhanced activity and explored the impacts of NAM on oligodendrocyte development and maturation.

Here, we demonstrate stage-specific exposure to NAM prior to the OPC expansion period significantly enhances neuronal activity in OCOs. The enhanced activity in OCOs was associated with increased organoid size, enhanced synaptic maturation, and heightened cellular heterogeneity as demonstrated by the emergence of diverse neuronal populations. Importantly, oligodendrocyte development and maturation are preserved in NAM conditions. Altogether, these studies establish an OCO model with enhanced neuronal maturation and activity while maintaining oligodendrocyte lineage progression in a heterogeneous, physiologically relevant cellular environment.

## Methods

### iPSC Culture

Four iPSC lines, consisting of three female and one male line, were included. Two female lines (WC-24-02-DS-B and ILD11-3) were thoroughly profiled and used in our previous studies (Campbell et al., 2023; Klein et al., 2022; Li et al., 2022). The (DS1-iPS4-disomic) male line was generated by Dr. Orkin’s lab (MacLean et al., 2012; Park et al., 2008) and recently acquired through Boston Children’s Hospital. An additional male control cell line (BU3-10-Cr2) was obtained from the Center of Regenerative Medicine (CReM), Boston University Chobanian & Avedisian School of Medicine. These lines are routinely maintained, tested for mycoplasma (monthly) and sterility and exhibit normal karyotypes. All iPSCs were used before passage 50 and maintained on Matrigel (Corning Incorporated, Corning, NY, United States, Cat#354277) with mTeSR plus (STEMCELL Technologies, Vancouver, BC, Canada, Cat#100-0276) and passaged with ReLeSR (STEMCELL Technologies, Vancouver, BC, Canada, Cat#05872).

### Generation of organoids

OCOs were derived from iPSCs as previously established (Madhavan et al., 2018) and demonstrated in our prior work (Campbell et al., 2023; Li et al., 2022). To generate OCOs, iPSCs between passages 24 – 50 were washed with PBS (Corning Incorporated, Corning, NY, United States, Cat#MT21040CV) and dissociated into a single cell suspension with Accutase (Thermo Fisher Scientific, Waltham, MA, United States, Cat#A1110501) for ∼5 min. Dissociated cells were plated at a density of 1.5 x 10^4^ cells per well in 150µL mTeSR plus with 50 µM Y-27632 (Tocris Bioscience, Bristol, United Kingdom, Cat#12-541-0) in low adherence V-bottom 96 well plates (Corning Incorporated, Corning, NY, United States, Cat#7007). On day 1, 100µL of media was replaced with TeSR-E6 (STEMCELL Technologies, Vancouver, BC, Canada, Cat#05946), supplemented with 2.5µM Dorsomorphin (STEMCELL Technologies, Vancouver, BC, Canada, Cat#72102) and 10 µM SB-431542 (STEMCELL Technologies, Vancouver, BC, Canada, Cat#72234). On days 3 - 6, 100µL of media changes with TeSR-E6 with Dorsomorphin and SB-431542 were performed daily. On day 7, media was changed to Neurobasal A medium (Thermo Fisher Scientific, Waltham, MA, United States, Cat#10888022) supplemented with B-27 supplement minus vitamin A (Thermo Fisher Scientific, Waltham, MA, United States, Cat#12587010), GlutaMAX (Thermo Fisher Scientific, Waltham, MA, United States, Cat#35050061), 100 U/mL Penicillin/Streptomycin (Thermo Fisher Scientific, Waltham, MA, United States, Cat#15140122) and Primocin (Invivogen, San Diego, CA, United States, Cat#ant-pm-1). From days 7 - 25, media was supplemented with 20 ng/mL fibroblast growth factor 2 (FGF2; STEMCELL Technologies, Vancouver, BC, Canada, Cat#78003.2) and 20 ng/mL epidermal growth factor (EGF; STEMCELL Technologies, Vancouver, BC, Canada, Cat#78006.1). From days 7-15 half media changes were performed daily, and from 17-25 media was changed every other day. Between days 16 and 20, organoids were transferred to ultra-low attachment 24 well plates (Corning Incorporated, Corning, NY, United States, Cat#07200602) and cultured individually with one per well. From days 27 – 39, OCOs were maintained in Neurobasal A medium supplemented with B27 serum substitute minus vitamin A, GlutaMax, 100 U/mL Penicillin/Streptomycin, Primocin, brain-derived neurotrophic factor (BDNF; STEMCELL Technologies, Vancouver, BC, Canada, Cat#78005.1), 20 ng/mL neurotrophic factor 3 (NT-3; STEMCELL Technologies, Vancouver, BC, Canada, Cat#78074.1) and 1% Geltrex (Thermo Fisher Scientific, Waltham, MA, United States, Cat#A1569601). Between days 40-49, organoids were maintained in Neurobasal-A supplemented with B27 without vitamin A, GlutaMax, Penicillin/Streptomycin, and Geltrex. Half changes of media were performed every other day. To expand the population of glial progenitors and promote OPCs expansion and differentiation as described in (Madhavan et al., 2018), 10 ng/mL platelet-derived growth factor AA (PDGF-AA; R&D Systems, Minneapolis, MN, United States, Cat#AF-307-SP) and 10 ng/mL insulin-like growth factor (IGF; Peprotech, Cranbury, NJ, United States, Cat#100-11) were added with fresh media every two days. From days 61 - 69, 40 ng/mL 3,3′,5-triiodothyronine (T3; Sigma-Aldrich, St. Louis, MO, United States, Cat#T6397) was added every two days. From day 70 onwards, 1 mL media changes were performed every 2 days. OCOs were transitioned to BrainPhys at days 40 and 70 or maintained in Neurobasal-A with supplements depending on the treatment protocol.

COs were generated as described in (Paşca et al., 2015), following the same protocol as OCOs with the exception of PDGF-AA, IGF, and T3.

NAM was composed of BrainPhys Neuronal Medium (STEMCELL Technologies, Vancouver, BC, Canada, Cat#05790) and supplemented with B-27 (Thermo Fisher Scientific, Waltham, MA, United States, Cat#17504001), 1% Matrigel, N-2 Supplement (STEMCELL Technologies, Vancouver, BC, Canada, Cat#07152), 10mM D-Glucose (Sigma-Aldrich, St. Louis, MO, United States, Cat#G8270), Lipid Concentrate (Thermo Fisher Scientific, Waltham, MA, United States, Cat#11905031), L-Ascorbic Acid (STEMCELL Technologies, Vancouver, BC, Canada, Cat#100-1040), 100 U/mL Penicillin/Streptomycin and Primocin. OCOs were gradually transitioned to NAM by increasing the composition by 20% each media change over the course of 10 days. After reaching 100% NAM, 1 mL half changes were performed every 2 days.

### Cryoprotection and Immunohistochemistry (IHC)

OCOs were fixed in 4% paraformaldehyde overnight, washed in PBS, and stored in 30% sucrose (Sigma-Aldrich, St. Louis, MO, United States, Cat#S0389). Organoids were embedded in a solution of optimum cutting temperature (OCT) compound (Fisher Scientific, Waltham, MA, United States, Cat#23-730-571) and 30% sucrose at a 60:40 ratio, flash frozen for storage at -80C and cryosectioned into 12µM sections.

For IHC, sections were permeabilized in PBS with 0.25% Triton (Sigma-Aldrich, St. Louis, MO, United States, Cat#X100) and blocked for 1 hour in 3% donkey serum (Sigma-Aldrich, St. Louis, MO, United States, Cat#D9663). Primary antibodies were diluted in blocking solution and incubated overnight at 4C. Sections were washed three times with PBST before application of secondary antibodies for 1 hour at room temperature. Slides were cover slipped with ProLong Gold Antifade Mountant with DAPI (Thermo Fisher Scientific, Waltham, MA, United States, Cat#P36931). The following primary antibodies were used: mouse CC-1 (1:250, Sigma-Aldrich (Calbiochem), St. Louis, MO, United States, Cat#OP80), goat SOX2 (1:200, R&D Systems, Minneapolic, MN, United States, Cat#AF2018), rat MBP (1:200, Abcam, Cambridge, United Kingdom, Cat#ab7349), mouse SOX2 (1:250, Abcam, Cambridge, United Kingdom, Cat#ab79351), rabbit PSD-95 (1:500, Invitrogen, Carlsbad, CA, United States, Cat#51-6900), mouse Synaptophysin (1:500, Santa Cruz Biotechnology, Dallas, TX, United States, Cat#sc-17750), guinea pig MAP2 (1:500, Synaptic Systems, Gottingen, Germany, Cat#188 004), mouse VGAT (1:400, Synaptic Systems, Gottingen, Germany, Cat#131 011), rat CTIP2 (1:100, Abcam, Cambridge, United Kingdom, Cat#ab18465), mouse SATB2 (1:500, Abcam, Cambridge, United Kingdom, Cat#ab51502), and goat Reelin (1:50, Thermo Fisher Scientific, Waltham, MA, United States, Cat#PA5-43537). The following secondary antibodies were used at a dilution of 1:500 to visualize primary antibodies: anti-mouse 546 (Thermo Fisher Scientific, Waltham, MA, United States, Cat#A11055), anti-goat 488 (Thermo Fisher Scientific, Waltham, MA, United States, Cat#A11039), anti-rat 647 (Jackson ImmunoResearch Laboratories, West Grove, PA, United States, Cat#712-605-153), anti-guinea pig 405 (Jackson ImmunoResearch Laboratories, West Grove, PA, United States, Cat#706-475-148) and anti-rabbit 750 (Abcam, Cambridge, United Kingdom, Cat#ab175728). Three sections per organoid were imaged on an Olympus VS200 Slide Scanner with a 20x objective. The total number of cells positive for a particular marker were quantified with QuPath (Bankhead et al., 2017). Representative whole organoid sections were imaged on a BC43 spinning disk confocal microscope (Oxford Instruments, High Wycombe, United Kingdom).

### Synaptic Marker Analysis

A 40X/NA1.3 oil objective on a BC43 spinning disk confocal microscope was used to image four separate fields of view with a z-step size of 0.1µm across two sections per organoid. PSD-95, synaptophysin and VGAT were quantified with Imaris v10.2.0 (Bitplane, Belfast, United Kingdom). Neuronal processes were defined by rendering a ‘Surface’ representation in Imaris and applying a manual threshold with background subtraction and smoothing enabled. Synaptic markers were detected using the ‘Spots’ tool and filtered using the ‘Quality’ parameter with Imaris machine learning segmentation. Colocalization was defined as PSD-95 puncta less than 1µm apart from Synaptophysin and within 1.5µm of MAP2, and puncta were normalized to MAP2 surface area.

### Organoid Area Measurement

OCOs were imaged under brightfield light microscope at indicated time points. Area was quantified by tracing organoid outlines on Fiji (Schindelin et al., 2012).

### Calcium Imaging

OCOs were transduced with 0.5µL of pGP-AAV-syn-jGCaMP8s-WPRE (Addgene, Watertown, MA, United States, Cat#162374) ∼ 2 weeks prior to imaging. For viral transduction, OCOs were incubated with ∼300µL of AAV-containing media overnight and placed back in the low attachment 24 well plates the following day. On the day of imaging, OCOs were transferred to Neurobasal-A minus phenol red (Thermo Fisher Scientific, Waltham, MA, United States, Cat#12349015) or BrainPhys imaging optimized medium (STEMCELL Technologies, Vancouver, BC, Canada, Cat#05796) with corresponding supplements according to culture condition.

A Zeiss LSM 710-Live Duo Confocal with a 10X objective was used to image GCaMP8s-labeled neurons. GCaMP8s was excited with a 488-nm laser and captured in a single z-plane for 180 seconds at a rate of 1 frame per second. OCOs were incubated at 37C and 5% CO2 throughout imaging and each organoid was captured across three consecutive fields of view.

### Calcium Imaging Analysis

Cytoplasmic calcium signals were extracted through manual region of interest (ROI) detection in Fiji. One background ROI excluding signal from nearby cells was subtracted from each video and three videos per organoid were analyzed. Raw traces were normalized to their baseline, defined as the mean of the lowest 15% of fluorescence intensity values (F0). To reduce noise, data was smoothed over a three-frame sliding window. Custom MATLAB (MathWorks, Natick, MA, United States, 2025b) scripts were used to perform analysis of calcium activity. Calcium transients were identified with ‘MinPeakProminence’ argument of the *findpeaks* function with a threshold of 0.5. A neuron was defined as active if at least one calcium transient was detected during the imaging period.

### OCO dissociation and scRNA-seq Library Preparation

On day 120, OCOs from the WC-24-02-DS-B cell line were dissociated with a papain dissociation system (Worthington Biochemical Corp., Lakewood, NJ, United States, Cat#LK003150) as previously described (Li et al., 2022). For each sample, three organoids from 2 independent differentiations were pooled, cut into small pieces, and dissociated in 0.005% DNase solution and 20 units/mL papain solution with 40 minutes of constant agitation. The papain solution was oxygenated with 95% O2 and 5% CO2. The cell suspension was then titrated with a 5mL pipette and centrifuged at 300 g for 2 minutes at room temperature and resuspended in PBS + 1% BSA. Samples with at least 88% viability as assessed by trypan blue staining were collected for target capture of 4,000 cells with 10X Genomics Chromium single cell preparation system according to manufacturer instructions.

cDNA synthesis, amplification, and library preparation were performed with 10X Genomics Chromium Single Cell 3’ Library and Bead Kit (v3) and sequenced on NovaSeq 6000 at the Single Cell Sequencing Core at Boston University School of Medicine.

### scRNA-seq Analysis

Raw Illumina BCL files were demultiplexed with bcl2fastq (v2.20). The count matrix was generated by aligning reads to the GRCh38 human reference genome (hg38) with 10X Genomics Cell Ranger (v9.0.1) and imported to Seurat (v5.1.0; (Hao et al., 2021) for downstream analysis. Cells with 200 – 10,000 unique genes and less than 15% mitochondrial reads were retained. Genes expressed in fewer than three cells were discarded. Doublets were removed with scDblFinder (v1.12.0; Germain et al., 2022). Datasets were aligned using ‘IntegrateLayers’ with canonical correlation analysis (CCA) integration.

Following integration, the data was log normalized and the top 2,000 highly variable features were identified using the VST method. The resulting features were scaled prior to performing dimensionality reduction with principal component analysis (PCA). The top 50 principal components (PCs) were selected and used to construct a shared nearest-neighbor (SNN) graph using ‘FindNeighbors’ followed by Louvain clustering with ‘FindClusters’ (resolution = 0.5). The data was then visualized with a uniform manifold approximation and projection (UMAP) plot. Clusters were manually annotated based on the expression of canonical marker genes and cluster-specific differentially expressed genes (DEGs). ‘FindMarkers’ was used to identify cluster markers with a Wilcoxon rank-sum test.

For DEG analysis, the ExN clusters were grouped based on layer identity. DEGs for the individual and grouped clusters were identified with a MAST test using ‘FindMarkers’. ClusterProfiler (v4.14.6) (Yu et al., 2012) was used to identify Gene Ontology (GO) terms from DEGs with a Benjamini-Hochberg adjusted p-value < 0.01 and log fold change greater than 0.25. The 29,994 genes expressed in the dataset were submitted as a background list. GO terms with a Benjamini-Hochberg adjusted p-value < 0.05 were considered significantly enriched. To reduce redundant GO terms, the simplify() function from ClusterProfiler with a cutoff of 0.7 was applied.

### Statistics

Data is expressed as mean ± SEM. Statistical significance was assessed using a two-tailed Welch’s t-test when two groups are compared. For the calcium imaging, a Mann-Whitney test was used when comparing two groups, and a Kruskal-Wallis test followed by Dunn’s multiple comparison was used when comparing multiple groups. A two-way ANOVA with Tukey post-hoc testing was used to evaluate significance for the area measurements. Statistical significance was indicated as: p > 0.05: ns, * p ≤ 0.05, ** p ≤ 0.01, *** p ≤ 0.001, **** p ≤ 0.0001. Statistical analysis was performed with GraphPad Prism 10.

## Results

### Exposure to NAM at day 40 maximally enhances activity in OCOs

To date, most studies utilizing organoids have focused on oligodendrocyte biology and functional neuronal readouts in isolation. To establish baseline levels of neuronal activity in oligodendrocyte-enriched cortical organoids (OCOs), we first generated organoids patterning the dorsal forebrain from one iPSC line with normal karyotype (**Figure S1**; Paşca et al., 2015) and expanded a small, native population of glial progenitors following neuronal expansion as previously described (**Figure 1A**; Madhavan et al., 2018) to generate OCOs. To evaluate the degree of measurable spontaneous neuronal activity in OCOs maintained NM, calcium imaging was performed in OCOs at day 110. Neurons were labeled with an adeno-associated virus (AAV) encoding GCaMP8s ∼2 weeks prior to imaging (**Figure 1B**) and defined as active if at least one calcium transient was detected over the 180s recording period. We observed minimal spontaneous activity in OCOs maintained in NM, with only 2.67% of neurons displaying detectable calcium activity (**Figure 1C**). We then validated this finding in an additional three iPSC lines and observed minimal spontaneous activity across independent cell lines (**Figure S2).** Following this observation, we aimed to develop a strategy for enhancing neuronal activity in OCOs while preserving the development of oligodendrocyte lineage cells.

**Figure 1.**
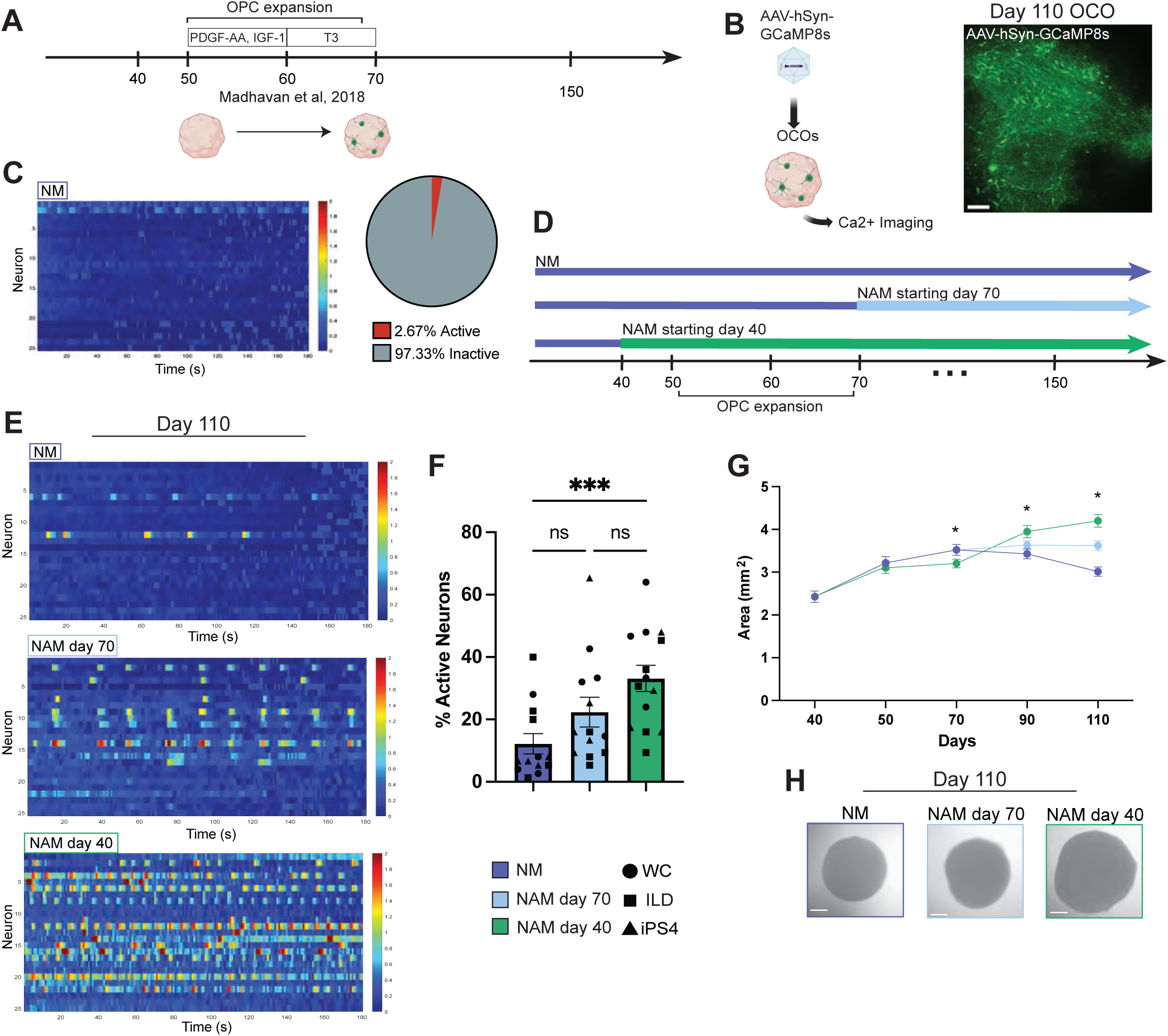
Exposure to NAM prior to OPC expansion enhances neuronal activity and OCO size. (A) Timeline for generation of oligodendrocyte-enriched cortical organoids (OCOs) as described in Madhavan et al. Graphics created with BioRender.com (B) Experimental workflow for functional assessment of OCOs using calcium imaging and representative image of GCaMP8s-labeled neurons. Scale bar is 100µm. Graphics created with Biorender.com. (C) Representative calcium imaging heatmap with ΔF/F traces of NM-OCOs. Pie chart displays the proportion of active neurons in NM-OCOs from BU3 iPSC line. N = 5 OCOs per experimental condition. (D) Schematic for the exposure to NAM in OCOs before (day 40) and after the OPC expansion period (day 70). (NM, Neuronal Medium; NAM, Neuronal Activity Medium). (E) Representative calcium imaging heatmaps of ΔF/F traces over 180s for each condition. (F) Percentage of active neurons in OCOs. Neurons are considered active if at least one calcium transient is detected in the 180s recording period. Each data point represents one OCO, averaged across three separate fields of view (FOV). N = 13 – 14 OCOs per experimental condition, collected from 3 independent iPSC lines. **** p ≤ 0.0001, Kruskal-Wallis test with Dunn’s multiple comparison. Error bars represent mean ± SEM. (G)Longitudinal area measurements of OCOs from days 40 – 110. n = 22 – 38 OCOs per timepoint and per experimental condition, collected from 3 independent iPSC lines. * p ≤ 0.05 between NAM day 40 and NM. Two-way ANOVA followed by Tukey’s post-hoc multiple comparisons test. Error bars represent mean ± SEM. (H) Representative brightfield images of day 110 OCOs in three different experimental conditions. Scale bar is 50µm.

Conventional basal media does not support the spontaneous firing of neurons (Bardy et al., 2015). To overcome this limitation in OCOs, we used NAM, a media permissive to spontaneous neuronal activity in vitro and aimed to establish the optimal developmental window where exposure to NAM would yield the most robust increase in neuronal activity.

To generate OCOs, OPC expansion and differentiation occurs through exposure to PDGF-AA and IGF from days 50 – 60, followed by T3 from days 60 – 70 (**Figure 1A**; Madhavan et al., 2018). Based on this framework, we incorporated NAM at two key timepoints: prior to the OPC expansion period (before day 50) and after OPC expansion and differentiation (day 70). In parallel, a control group of OCOs was maintained in NM, (hereafter referred to as NM-OCOs; **Figure 1D**).

Before introducing NAM, we first confirmed that neural progenitor composition was comparable across cell lines to ensure a consistent baseline prior to intervention. At day 40, preceding both media manipulation and OPC expansion, the proportion of SOX2+ neural progenitors and SOX2+/Ki67+ cycling progenitors were similar across four cell lines (**Figure S3A, B**). As expected, SOX10 expression was minimal at this stage, consistent with the absence of oligodendrocytes prior to the OPC expansion phase (Madhavan et al., 2018; **Figure S3A, B**).

Following the experimental design outlined in Figure 1D, we gradually transitioned OCOs to NAM over a 10-day period beginning at either day 40 or 70. Initiating NAM exposure at day 70 resulted in a modest and nonsignificant increase in spontaneous activity relative to NM-OCOs (22.36% active day 70 NAM-OCOs vs. 12.2% active NM-OCOs; p = 0.1765; **Figure 1E, F**). In contrast, OCOs transitioned to NAM starting at day 40, and thus exposed to NAM prior to OPC expansion, displayed a robust increase in activity (33.14% active neurons in day 40 NAM-OCOs vs. 12.2% in NM-OCOs; p = 0.0008; **Figure 1E, F**). Thus, early NAM exposure, preceding the OPC expansion phase, appears to maximally enhance spontaneous neuronal activity in OCOs.

We next evaluated the impacts of exposure to NAM on cortical organoids (COs) without oligodendrocyte enrichment as described in (Paşca et al., 2015). Similar to OCOs, we observed minimal neuronal activity in NM-COs and a significant increase in activity following introduction to NAM at day 40 (9.56% active NM-COs vs. 28.67% active day 40 NAM-COs, p = 0.0416; **Figure S4A, B**). However, in contrast to the OCOs, neuronal activity was most significantly increased following exposure to NAM at day 70 (9.56% active NM-COs vs. 34.91% active NM-COs, p = 0.0009; **Figure S4A, B**). Thus, exposure to NAM at both day 40 and day 70 significantly enhances activity in COs, while exposure to NAM at day 40, prior the OPC expansion period, is necessary to enhance activity in OCOs.

In addition to our functional assessments, OCO area was quantified beginning at day 40. Initially, OCOs exposed to NAM at day 40 showed a small but significant decrease in area at day 70 compared to NM-OCOs (3.10mm^2^ day 40 NAM-OCOs vs. 3.22mm^2^ NM-OCOs; p = 0.0188; **Figure 1G**). However, longitudinal measurements revealed a consistent and significant increase in day 40 NAM-OCO size relative to NM controls at day 90 (3.63 mm^2^ day 40 NAM-OCO vs. 3.43 mm^2^ NM-OCOs; p = 0.0188; **Figure 1G**) and day 110 (4.20 mm^2^ day 40 NAM-OCOs vs. 3.01 mm^2^ NM-OCOs; p = 0.0188; **Figure 1G, H**). In contrast, NAM-OCOs exposed to NAM beginning at day 70, did not display a significant difference in size relative to NM-OCOs at these later time points (**Figure 1G**). Notably, similar to NM-OCOs, day 70 NAM-OCOs stopped increasing in size over the culture duration (**Figure 1G**).

Similarly to OCOs, COs treated with NAM beginning at day 40 display a significant reduction in size relative to NM-COs beginning at days 50 (3.10 mm^2^ day 40 NAM-OCOs vs. 3.62 mm^2^ NM-OCOs; p = 0.0422; **Figure S4C**) and 70 (3.72 mm^2^ day 40 NAM-OCOs vs. 3.96 mm^2^ NM-OCOs; p = 0.0422; **Figure S4C**), followed by a significant increase in size on days 90 (4.91 mm^2^ day 40 NAM-OCOs vs. 3.82 mm^2^ NM-OCOs; p = 0.0422; **Figure S4C**) and 110 (5.27 mm^2^ day 40 NAM-OCOs vs. 3.27 mm^2^ NM-OCOs; p = 0.0422; **Figure S4C, D**). Meanwhile, COs maintained in NM and those transition to NM on day 70 displayed minimal expansion past day 50.

Together, organoid size in both COs and OCOs corresponded with the degree of spontaneous activity (**Figure 1F**, **Figure S4**), which may reflect enhanced neuronal growth and branching under the more active conditions. As OCOs are the primary focus of this study, we prioritized this system for subsequent analyses. Based on the coordinated increase in neuronal activity and organoid size, we defined day 40 as the optimal time window for NAM exposure and focused all subsequent analyses on OCOs exposed to NAM at this timepoint (hereafter NAM-OCOs).

### Cellular heterogeneity is increased in NAM-OCOs

To link our functional findings with compositional and molecular scale changes, we performed transcriptomic profiling of NM and NAM-OCOs at day 110. NM and NAM-OCOs collected from two independent differentiations were pooled together for single cell RNA-sequencing (scRNA-seq). Following quality control filtering, 8,933 cells were retained, yielding 17 transcriptionally distinct clusters (**Figure 2A**). Annotation of cluster-specific specific differentially expressed genes (DEGs) and canonical marker genes resulted in the identification of seven major cell types, encompassing excitatory neurons (ExN), inhibitory neurons (InN), radial glia cells (RGC), intermediate progenitor cells (IPC), Reelin+ cells (RELN), astrocytes (Astro), and oligodendrocyte lineage cells (OLLC) (**Figure 2A; Figure S5A, B**).

**Figure 2.**
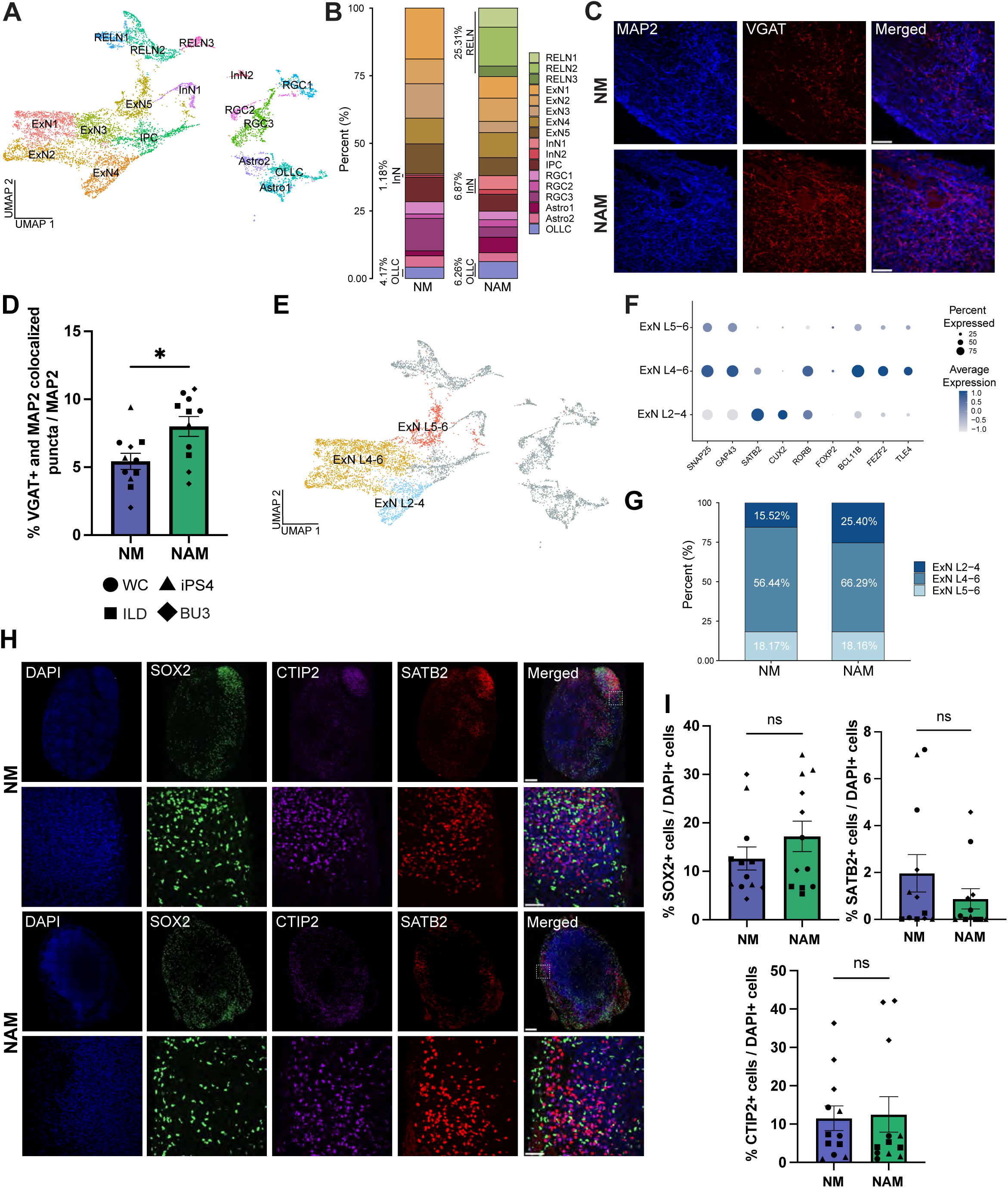
Cellular diversity is enhanced in day 110 NAM-OCOs. (A) UMAP plot of scRNAseq data from day 110 OCOs. RGC, radial glial cells; IPC, intermediate progenitors; ExN, excitatory neurons; InN, inhibitory neurons; RELN, Reelin+ cells; Astro, Astrocytes; OLLC, Oligodendrocyte lineage cells. 3-4 OCOs from 2 independent differentiation were used. (B) Stacked bar plot displaying the percent composition of NM and NAM-OCO clusters. (C) Representative immunofluorescence images of day 110 OCOs stained with MAP2 and VGAT. Scale bar is 50µm. (E) Quantification of the percentage of VGAT puncta normalized to total MAP2 using Imaris v10.2.0. Each data point represents one OCO, averaged across four fields of view from two separate sections, and each shape represents a separate cell line. N = 11 OCOs per experimental condition, collected from 4 independent iPSC lines. * p ≤ 0.05, two-tailed Welch’s t-test. Error bars represent mean ± SEM (F) UMAP highlighting ExN populations identified and used for differential expression analysis (G) Dot plot of ExN marker genes used to identify ExN subtypes. (H) Representative immunofluorescence images of day 110 OCOs stained with SATB2, CTIP2 and SOX2. Scale bar, 200µm and 50µm, inset. (I) Quantification of CTIP2, SATB2, and SOX2. Expression of markers is normalized to DAPI and quantified with QuPath v0.6.0. Each data point represents one OCO, averaged across three sections, and each shape represents a separate cell line. N = 12 OCOs per experimental condition, collected from 4 independent iPSC lines. ns = no significance, two-tailed Welch’s t-test. Error bars represent mean ± SEM

Our transcriptomic analysis revealed a shift in cellular composition in NAM-OCOs relative to NM-OCOs. Specifically, we noted increased proportions of GABAergic InNs and a slight increase in the proportion of OLLCs in NAM-OCOs (**Figure 2B**). NAM-OCOs were composed of 6.87% InNs and 6.26% OLLCs, relative to 1.18% and 4.17% in NM-OCOs, respectively (**Figure 2B**). We also discovered an enrichment in a population of Reelin+ cells in NAM-OCOs (**Figure 2B, Figure S5A, B**). To validate these findings across OCOs generated from independent iPSC lines, we performed immunostaining for Reelin in day 110 NM and NAM-OCOs (**Figure S6A)**. Quantification of Reelin+ cells revealed that they are present at low abundance in NM-OCOs and were likely not captured in our sequencing data. NAM-OCOs displayed a 54.44% increase in the proportions of Reelin+ cells, although this did not reach statistical significance (p = 0.0544; **Figure S6B**). These results suggest that early NAM exposure may enhance Reelin+ cells.

We next validated the increased InN population through immunostaining for the vesicular GABA transporter (VGAT) and microtubule-associated protein 2 (MAP2). In line with our transcriptomic findings, NAM-OCOs exhibited a significant increase in VGAT+ puncta/MAP2 (8.00% VGAT+ puncta/MAP2+ in NAM-OCOs vs. 5.43% in NM-OCOs, p = 0.0133; **Figure 2C, D**), indicating an expanded population of GABAergic InNs in NAM-OCOs. Together, these findings demonstrate that early NAM exposure alters cellular composition, promoting InN enrichment in OCOs.

We next investigated whether NAM impacts excitatory neuron production. Our transcriptomic analysis demonstrated comparable proportions of deep layer (ExN4-6 and ExN5-6) and superficial layer (ExN L2-4) neurons in NM-OCOs and NAM-OCOs (**Figure 2G)**. In line with this, immunostaining of deep and superficial layer neuronal markers revealed no significant differences in the percentage of CTIP2+ deep layer (12.51% CTIP2+ cells NAM-OCOs vs. 11.52% NM-

OCOs; p = 0.8618) or SATB2+ superficial layer neuronal markers (0.88% SATB2+ cells NAM-OCOs vs. 1.97% NM-OCOs; p = 0.2476). Further, both NM and NAM-OCOs maintained comparable pools of SOX2+ progenitors at day 110 (12.64% SOX2+ cells NM-OCOs vs. 17.23% NAM-OCOs; p = 0.2578; **Figure 2H, I**). Together, these results suggest the increased size of NAM-OCOs is likely not driven by an expanded progenitor pool or altered ExN production but may reflect potentially enhanced neurite branching and complexity.

### Structural maturation is enhanced in NAM-OCOs

Following our observation of preserved ExN composition, we examined whether NAM is associated with transcriptional changes by performing differential expression analysis within our ExN subpopulations (**Figure 2E, F**). Analysis of upregulated DEGs following NAM exposure across upper-layer (ExN L2–3) and deep-layer neurons (ExN L4–6 and L5–6) revealed significant enrichment of biological processes related to mRNA processing, dendrite development, neuron projection organization, and synaptic transmission (**Figure 3A)**. In contrast, the downregulated DEGs were primarily enriched for terms related to cellular respiration, oxidative phosphorylation, mitochondrial organization, and cytoplasmic translation without enrichment in terms directly associated with neuronal differentiation or neurite organization (**Figure 3B**). This pattern was consistent across ExN layers, pointing to a coordinated global shift toward structural maturation and neuronal development in NAM-OCOs rather than a layer-specific effect.

**Figure 3.**
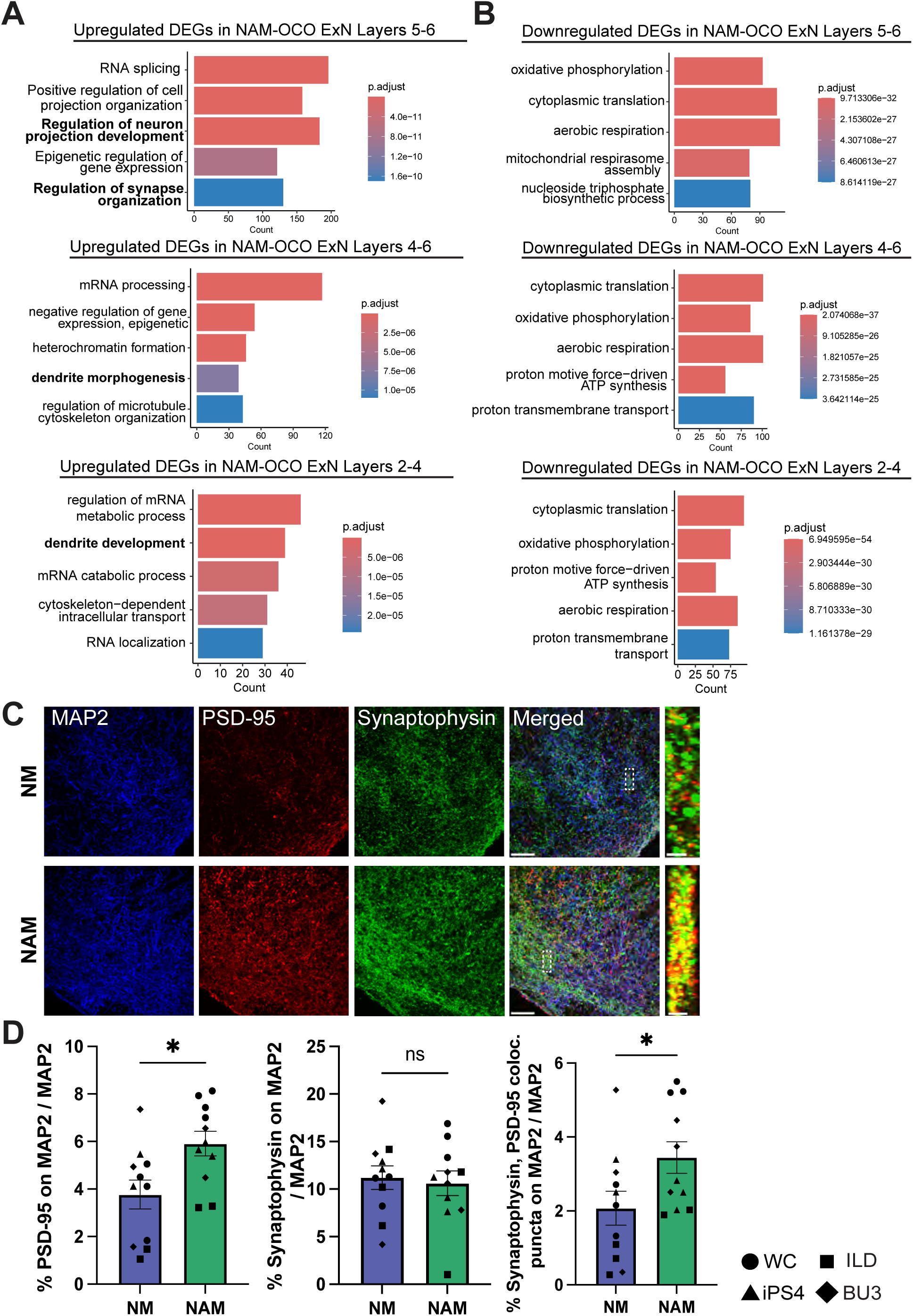
Structural maturation is enhanced in day 110 NAM-OCOs. (A-B) Enriched GO biological process (BP) terms of upregulated (A) and downregulated (B) differentially expressed genes from ExN layers 5-6, 4-6, and 2-4. (C) Representative immunofluorescence images and 3D cross-section of day 110 OCOs stained with MAP2, PSD-95 and Synaptophysin. Scale bar is 50µm and 2µm on cross-section. (D) Quantification of the percentage of Synaptophysin and PSD-95 colocalized puncta on MAP2, normalized to total MAP2, using Imaris v10.2.0. Each data point represents one OCO, averaged across four fields of view from two separate sections, and each shape represents a separate cell line. N = 11 OCOs per experimental condition, collected from 4 independent iPSC lines. * p ≤ 0.05, two-tailed Welch’s t-test. Error bars represent mean ± SEM.

To validate our findings of elevated expression of synaptic development genes in NAM-OCOs (**Figure 3A)**, we quantified synaptic density by evaluating colocalization of the excitatory post-synaptic density marker PSD-95 and the pre-synaptic marker synaptophysin along MAP2-positive neuronal soma and dendritic processes. Overlapping PSD-95 and synaptophysin puncta were defined as structurally assembled synapses. NAM-OCOs exhibited a significant increase in PSD-95 puncta density, (5.91% PSD-95+ puncta/MAP2 NAM-OCOs vs. 3.77% NM-OCOs; p = 0.0146; **Figure 3C, D**) and an overall increase in colocalization of pre- and post-synaptic markers (3.45% colocalized puncta/MAP2 NAM-OCOs vs. 2.07% NM-OCOs; p = 0.0408; **Figure 3C, D**). Thus, NAM-OCOs exhibit increased synaptic density, driven in part by a prominent increase in excitatory postsynaptic structures. This increase in PSD95+ puncta is consistent with the previously established role of post-synaptic density formation in synaptogenesis and synaptic stabilization (Cane et al., 2014; Nikonenko et al., 2008).

### Oligodendrocyte development is preserved in NAM-OCOs

To confirm enhancing neuronal activity does not come at the expense of oligodendrocyte lineage cells, we next investigated the impact of NAM on oligodendrocyte development and maturation We first assessed early oligodendrocyte differentiation by quantifying expression of CC1, a pan-oligodendrocyte marker, following the OPC expansion period at day 70. The number of CC1+ oligodendrocytes did not differ between conditions (5.61% NAM-OCOs CC1+ cells vs. 5.83% NM-OCOs; p = 0.8295; **Figure 4A, B**), indicating that NAM exposure does not affect oligodendrocyte expansion at early stages.

**Figure 4.**
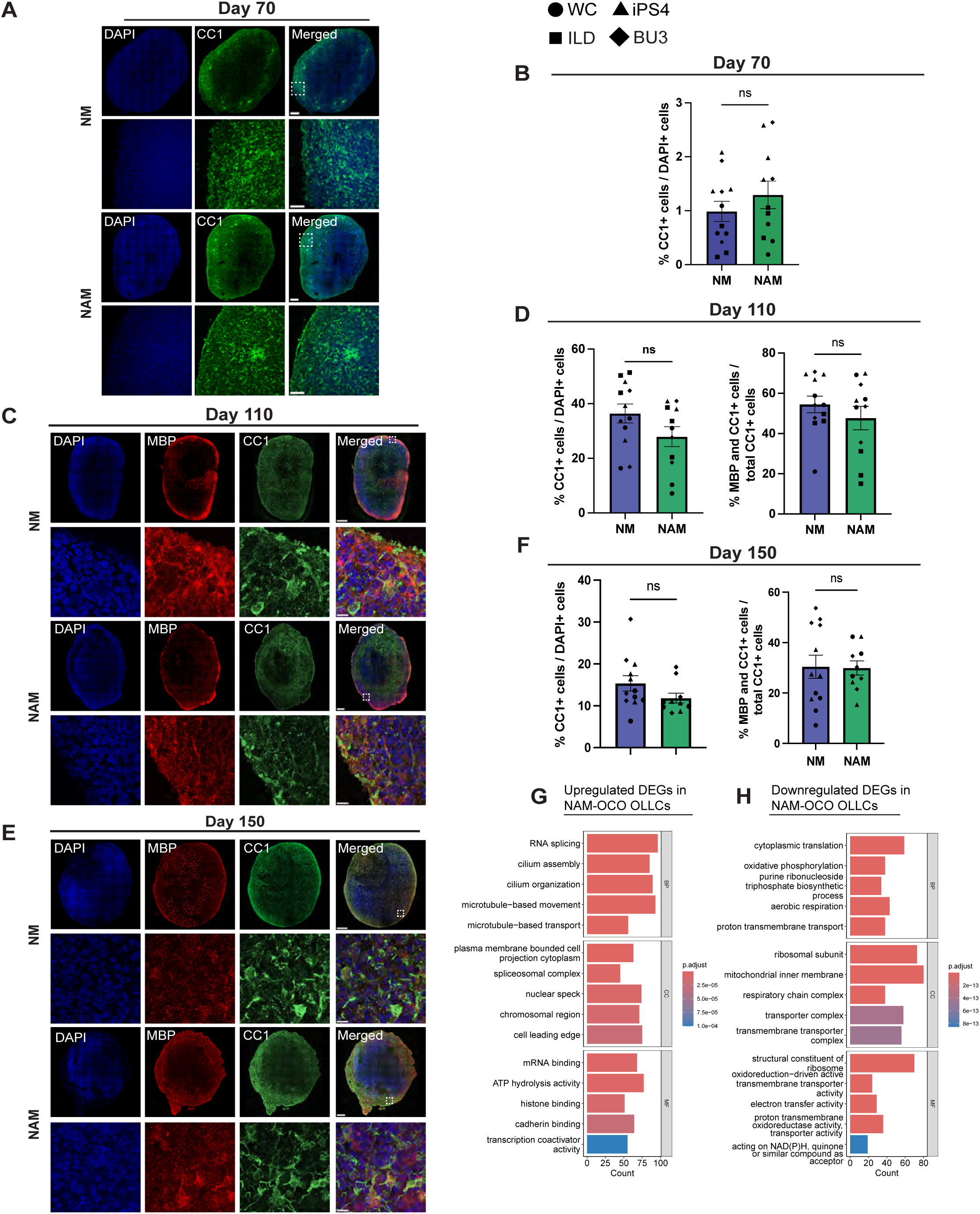
Oligodendrocyte development is preserved in NAM-OCOs. (A) Representative immunofluorescence images of day 70 OCOs stained with CC1. Scale bar, 200µm, 50µm inset. (B) Quantification of CC1. Expression of markers is normalized to DAPI and quantified with QuPath v0.6.0. Each data point represents one OCO, averaged across three sections, and each shape represents a separate cell line. N = 12 OCOs per experimental condition, collected from 4 independent iPSC lines. ns = no significance, two-tailed Welch’s t-test. Error bars represent mean ± SEM. (C – F) Representative immunofluorescence images of day 110 (C) and day 150 (E) OCOs stained with MBP and CC1. Quantification of total CC1+ cells normalized to DAPI+ cells, and the proportion of cells co-expressing MBP and CC1 normalized to the total number of CC1+ cells in day 110 (D) and day 150 (F) OCOs. Each data point represents one OCO, averaged across three sections, and each shape represents a separate cell line. N = 12 OCOs per experimental condition, collected from 4 independent iPSC lines. * p ≤ 0.05, ns = no significance, two-tailed Welch’s t-test. Error bars represent mean ± SEM. (G, H) Top 5 enriched GO biological process (BP), cellular component (CC), and molecular function (MF) terms in genes upregulated (G) and downregulated (H) in NAM-OCO OLLCs.

We next evaluated oligodendrocyte maturation at day 110, the same timepoint at which calcium imaging was performed, to confirm enhancing neuronal activity does not come at the expense of oligodendrocytes. Immunostaining of CC1 and MBP was performed to quantify numbers of CC1+ oligodendrocyte lineage cells and myelinating oligodendrocytes (CC1 and MBP+). No significant differences in the total numbers of CC1+ cells (27.95% CC1+ cells NAM-OCOs vs. 36.43% NM-OCOs, p = 0.1089; **Figure 4C, D**) or the proportion of CC1+ cells co-expressing MBP+ (47.67% CC1 and MBP+ cells NAM-OCOs vs. 54.49% NM-OCOs p = 0.3516; **Figure 4C, D**) were detected at day 110. Similarly, at day 150, which corresponds to a more mature state oligodendrocyte stage within OCOs (Madhavan et al., 2018) no differences in CC1+ cells (11.83% CC1+ cells NAM-OCOs vs. 15.34% NM-OCOs; p = 0.1250; **Figure 4E, F**) or CC1+ cells co-expressing MBP (29.91% CC1 and MBP+ cells NAM-OCOs vs. 30.42% NM-OCOs; p = 0.9259; **Figure 4E, F**) were observed. Thus, despite our transcriptomic data suggesting a small expansion of OLLCs following the exposure to NAM, no impact on the production of oligodendrocytes or MBP expression was detected at a protein level.

Although immunostaining did not suggest changes in oligodendrocyte numbers or maturity, we further explored transcriptional changes in our oligodendrocyte population following exposure to NAM by identifying DEGs in our OLLC cluster from our scRNAseq data (**Figure S5A, B).** In OLLCs, exposure to NAM was associated with upregulation of pathways relating to microtubule-based movement and cell projections (**Figure 4G**), consistent with enhanced structural maturation and process extension. Further, we observed enrichment for terms relating to cilium assembly and organization in OLLCs (**Figure 4G**), in line with past work demonstrating primary cilia mediate OPC proliferation and differentiation (Falcón-Urrutia et al., 2015). OLLCs exposed to NAM also displayed downregulation of cytoplasmic translation, oxidative phosphorylation, aerobic respiration, and mitochondrial organization (**Figure 4H**), mirroring the metabolic shift observed in ExNs. Together, our data suggest prolonged culture in NAM supports oligodendrocyte lineage development and results in a transition across ExNs and OLLCs towards structural reorganization and decreased expression of genes associated with oxidative metabolism.

## Discussion

Modeling neuronal activity and oligodendrocyte development is critical to capturing the functional interactions present *in vivo.* Here, using four independent iPSC lines, we demonstrate that stage specific exposure to NAM enhances neuronal activity in OCOs, establishing a model that integrates neuronal maturation with oligodendrocyte lineage development.

To enhance neuronal activity in OCOs, we introduced NAM before and after the OPC expansion period described in (Madhavan et al., 2018) while maintaining a control group in a standard basal NM. We demonstrated exposure to NAM at day 40, prior to expansion of the oligodendrocyte lineage, resulted in the most robust increase in neuronal activity relative to OCOs maintained in NM, whereas exposure to NAM after the oligodendrocyte enrichment period at day 70 did not significantly enhance activity relative to NM-OCOs. Notably, NAM significantly enhanced neuronal activity in COs, which do not undergo OPC expansion, at both day 40 and day 70, indicating that NAM exposure consistently enhances neuronal activity across organoid systems. However, as OCOs are the focus of our study, we used this system to interrogate neuron–oligodendrocyte co-development. Together, our results suggest exposure to NAM prior to the OPC expansion period is optimal for supporting the co-development of neurons and oligodendrocytes within OCOs.

In addition to our functional analyses, we performed transcriptomic and immunohistochemical profiling of NM and NAM-OCOs. While we discovered exposure to NAM does not significantly impact deep and superficial layer neuron composition, NAM-OCOs displayed increased cellular heterogeneity. In particular, we noted heightened proportions of GABAergic interneurons. Previous work has demonstrated inhibitory signaling from GABAergic interneurons improves network maturation and complexity in organoids (Quadrato et al., 2017; Samarasinghe et al., 2021; Trujillo et al., 2019). Thus, the increased interneuron population in NAM-OCOs may reflect greater circuit refinement and contribute to an increasingly mature network phenotype.

During maturation, neurons upregulate cytoskeletal proteins and enhance their morphological complexity (Compagnucci et al., 2016; Di Bella et al., 2021). We discovered significantly elevated size of NAM-OCOs, without corresponding changes in progenitor and laminar composition, suggesting that increased size can be attributed to heightened neurite outgrowth and branching. Moreover, transcripts upregulated in the excitatory neurons of NAM-OCOs primarily reflected biological processes related to neuronal projection and cytoskeletal dynamics across layers. Our findings support the conclusions of previous studies describing increased neuronal branching in cultures maintained in BrainPhys (Faravelli et al., 2025; Faria-Pereira et al., 2022; Satir et al., 2020). This enhanced morphological complexity may promote connectivity, and in conjunction with the enhanced cellular diversity and synaptic density, contribute to elevated activity in NAM-OCOs.

In addition to morphological and synaptic changes, neuronal maturation is accompanied by increased reliance on oxidative phosphorylation (OxPhos) to meet growing energetic demands (Iwata et al., 2023; Zheng et al., 2016). Consistent with this, previous work has demonstrated that BrainPhys increases OxPhos utilization in mouse primary neuronal cultures (Faria-Pereira et al., 2022). Unexpectedly, we discovered a transcriptional profile across our excitatory neuron and oligodendrocyte populations indicative of reduced OxPhos in NAM-OCOs. One potential explanation is the metabolic constraints inherent to 3D organoid systems. As OCOs increase in size, limited oxygen diffusion can create a hypoxic environment in the center of organoids (Uzquiano et al., 2022), favoring glycolytic metabolism. Although we did not observe a direct enrichment of glycolysis-related pathways in our NAM-OCOs, it is possible the larger size of NAM-OCOs results in a greater proportion of hypoxic cells, resulting in a shift away from oxidative metabolism. Thus, while NAM-OCOs display elevated structural and functional maturation, their metabolic phenotype may reflect an adaptive response to oxygen and diffusion limitations inherent to 3D organoid cultures.

Finally, we demonstrate that oligodendrocyte development and maturation are preserved in NAM-OCOs. While our transcriptomics suggested increased proportions of oligodendrocytes in NAM-OCOs, we discovered no significant differences in numbers of oligodendrocytes or proportions of myelinating oligodendrocytes in NM-OCOs and NAM-OCOs across multiple developmental timepoints, indicating oligodendrocyte lineage progression is preserved in NAM-OCOs. Inspection of differentially expressed genes suggested enhanced OPC differentiation and branching in NAM-OCOs, as indicated by the elevated expression of transcripts associated with microtubule dynamics, process intention, and ciliary function. Primary cilia serve as key signaling hubs for extracellular cues, including regulation of oligodendrocyte proliferation and differentiation (Falcón-Urrutia et al., 2015). While we did not directly observe differences in numbers of oligodendrocytes at a protein level, the transcriptional changes associated with heightened ciliary function and cytoskeletal dynamics may facilitate process extension and differentiation, potentially enhancing oligodendrocyte function.

Capturing neuronal activity and oligodendrocyte lineage cells presents a valuable opportunity for investigating oligodendrocyte-neuron interactions in a human cellular system. Whether enhanced neuronal activity with NAM influences oligodendrocyte biology or, conversely, whether reciprocal interactions might modulate neuronal activity merits further investigation. Among many other non-canonical functions, OPCs receive direct synaptic input from neurons (Bergles et al., 2000), phagocytose synapses (Auguste et al., 2022; Buchanan et al., 2022), and respond to neuronal activity (Geraghty et al., 2019; Gibson et al., 2014). Detailed understanding of these interactions in human cellular systems is limited; NAM-OCOs present a powerful tool for understanding how enhanced neuronal activity may modify these OPC-neuron interactions and further our understanding of their disruption in disease. Future studies using neuronal activity blockade will be required to directly establish causative relationship between activity and oligodendrocyte maturation in our system.

Altogether, we establish an increasingly comprehensive OCO model containing oligodendrocyte lineage cells and neuronal activity in a mature, heterogeneous cellular environment. We demonstrate stage-specific exposure to NAM enhances the functional and structural maturation of OCOs while preserving oligodendrocyte lineage integrity and maturation. Generating an organoid with mature oligodendrocytes and active neurons provides a basis for further study of the precise signaling mechanisms and cellular cues shaping oligodendrocytes and their interactions with neurons in healthy and pathological contexts.

## Acknowledgments

We thank Dr. Yuriy Alekseyev and the members of the Boston University Chobanian & Avedisian School of Medicine Microarray and Sequencing Resource Core for their help and guidance with the sequencing experiments. We are grateful to NIH agencies for their support: NIH/NIA RF1AG088529(MPI: E. Zeldich and M. Thunemann), NIH/NINDS R21NS125469 (MPIs: E Zeldich and M. Medalla), NIH/NIA and R03NS126864 (PI: E. Zeldich).

## Declaration of generative AI and AI-assisted technologies in the writing process

During the preparation of this work the authors used in ChatGPT (OpenAI) to assist with language editing, proofreading, and improving clarity of the manuscript. After using this tool, the authors reviewed and edited the content as needed and take full responsibility for the content of the published article.

## Author contributions

CC, EKK, and NBC designed and performed all aspects of the cell culture experiments and interpreted the results. CC and EZ wrote the first drafts of the manuscript. EZ conceived the idea and oversaw the project, provided guidance, interpreted the results, and edited the manuscript. CC and JLO designed and performed bioinformatics analyses for scRNA-seq dataset. CC, GF, KP, and MK performed quantitative analysis of immunohistochemical staining and image acquisition. CVG assisted with analysis of calcium imaging data.

## Resource availability

Raw and processed scRNA-seq data generated in this study have been deposited in the NCBI Gene Expression Omnibus database (GEO). Source data and code used for analysis will be made available by authors upon request.

