## Supplementary Figures S1, S2, S3, S4, S5 for "Stage-specific exposure to an activity-permissive media enhances neuronal maturation in oligodendrocyte-enriched cortical organoids"

Figure S1

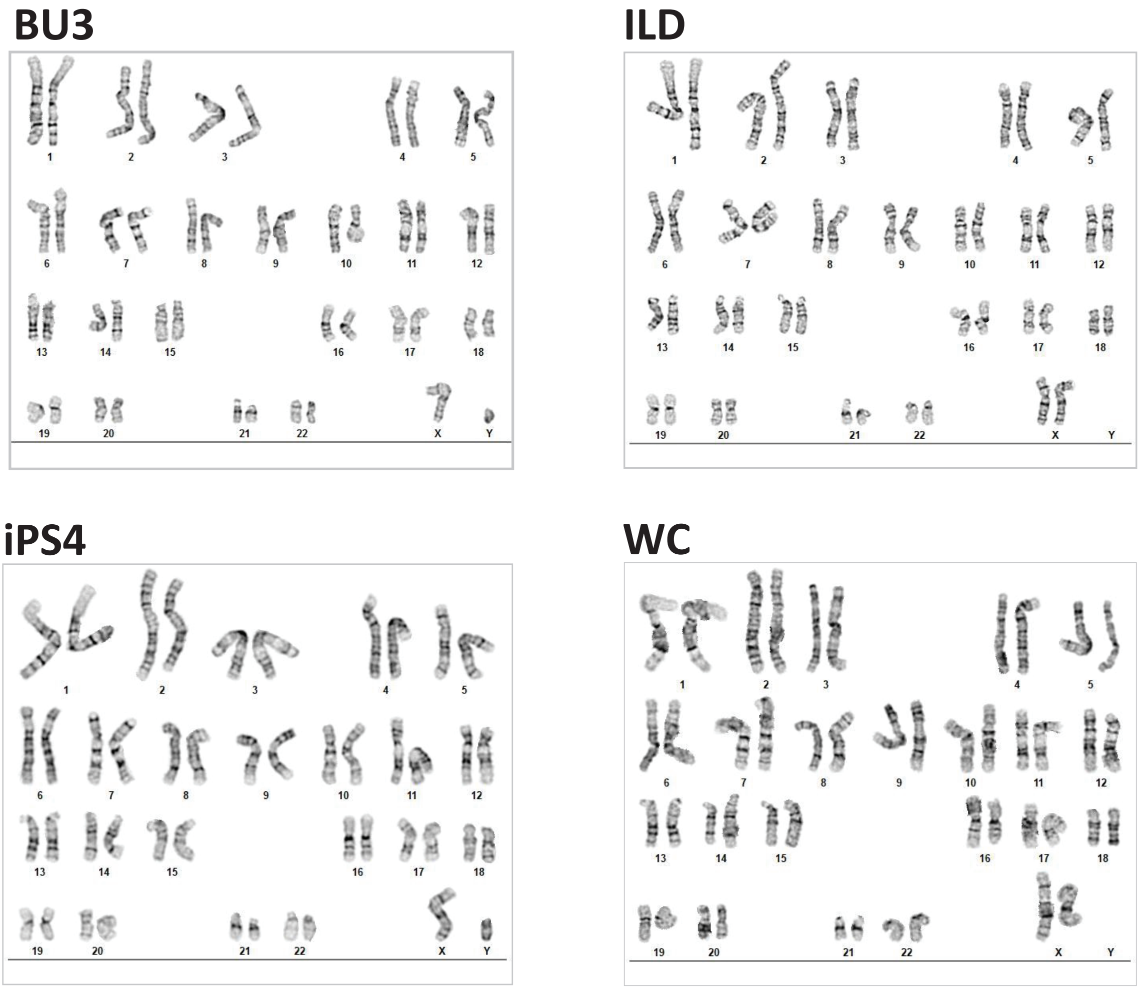
BU3
1 2 3 4 5
6 7 8 9 10 11 12
13 14 15 16 17 18
19 20 21 22 X Y
ILD
1 2 3 4 5
6 7 8 9 10 11 12
13 14 15 16 17 18
19 20 21 22 X Y
iPS4
1 2 3 4 5
6 7 8 9 10 11 12
13 14 15 16 17 18
19 20 21 22 X Y
WC
1 2 3 4 5
6 7 8 9 10 11 12
13 14 15 16 17 18
19 20 21 22 X Y

Figure S2

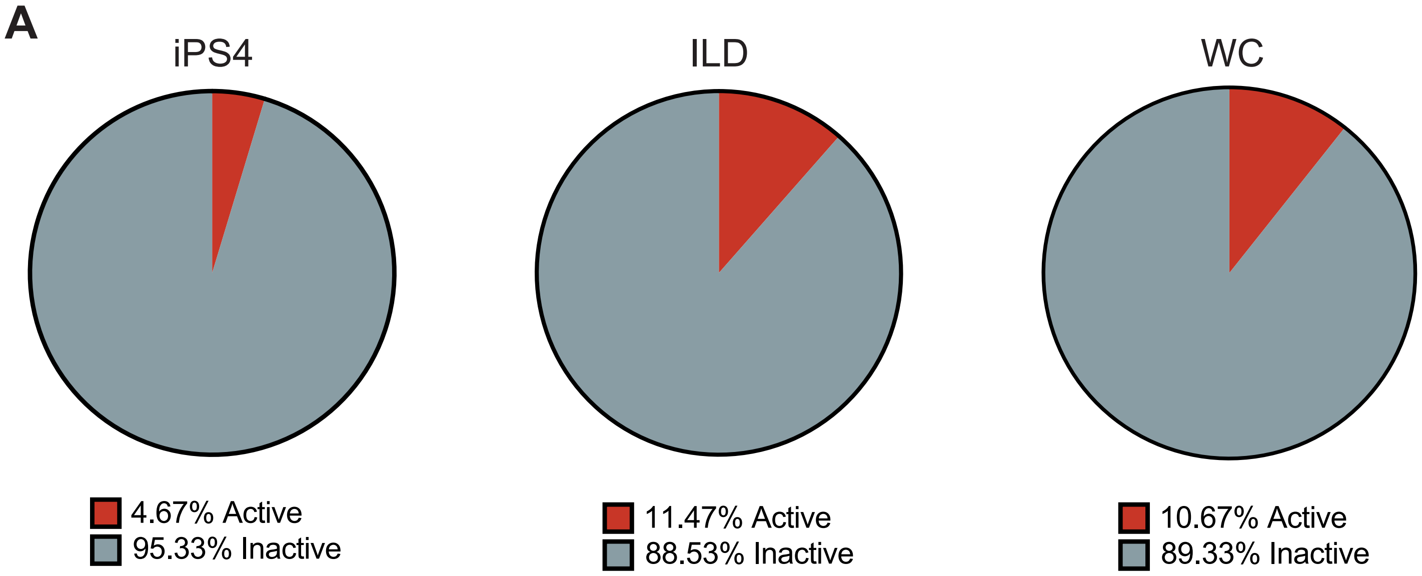


**Figure S2. Spontaneous activity levels in day 110 NM-OCOs.** (A) Pie charts depict the proportion of spontaneously active neurons in NM-OCOs derived from three iPSC lines. Neurons are considered active if at least one calcium transient is detected in the 180s recording period. N = 4 – 5 OCOs per line.

Figure S3

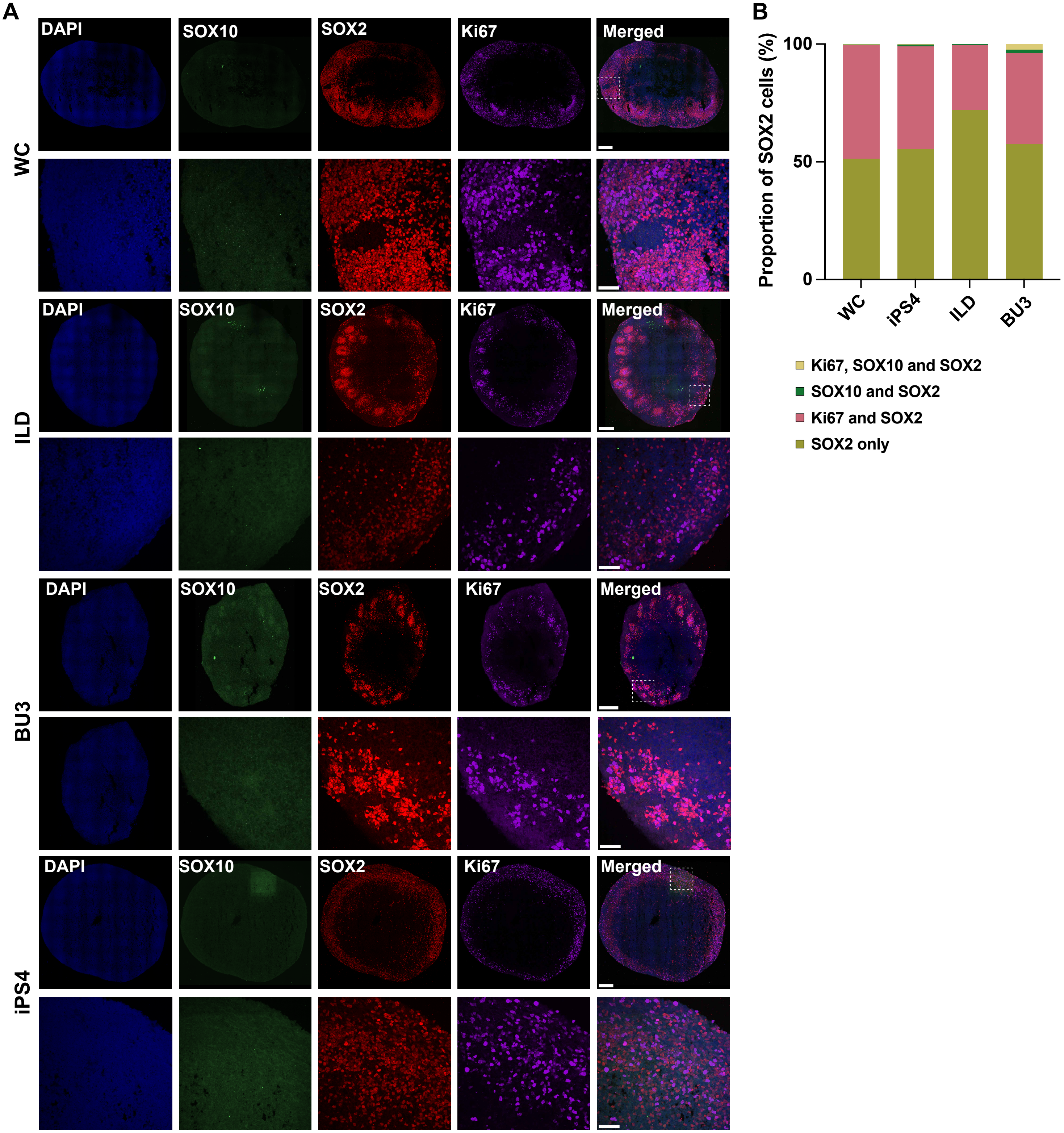


**Figure S3. Emergence of progenitors at day 40**. (A) Representative immunofluorescence images of day 40 organoids stained with SOX10, SOX2, and Ki67 from four iPSC lines. Scale bar = 200um and 50um, inset. (B) Relative proportions of SOX2+ progenitors co-expressing with SOX10, Ki67, or SOX10 and Ki67. Quantified with QuPath v0.6.0. N = 3 OCOs per cell iPSC line.

Figure S4

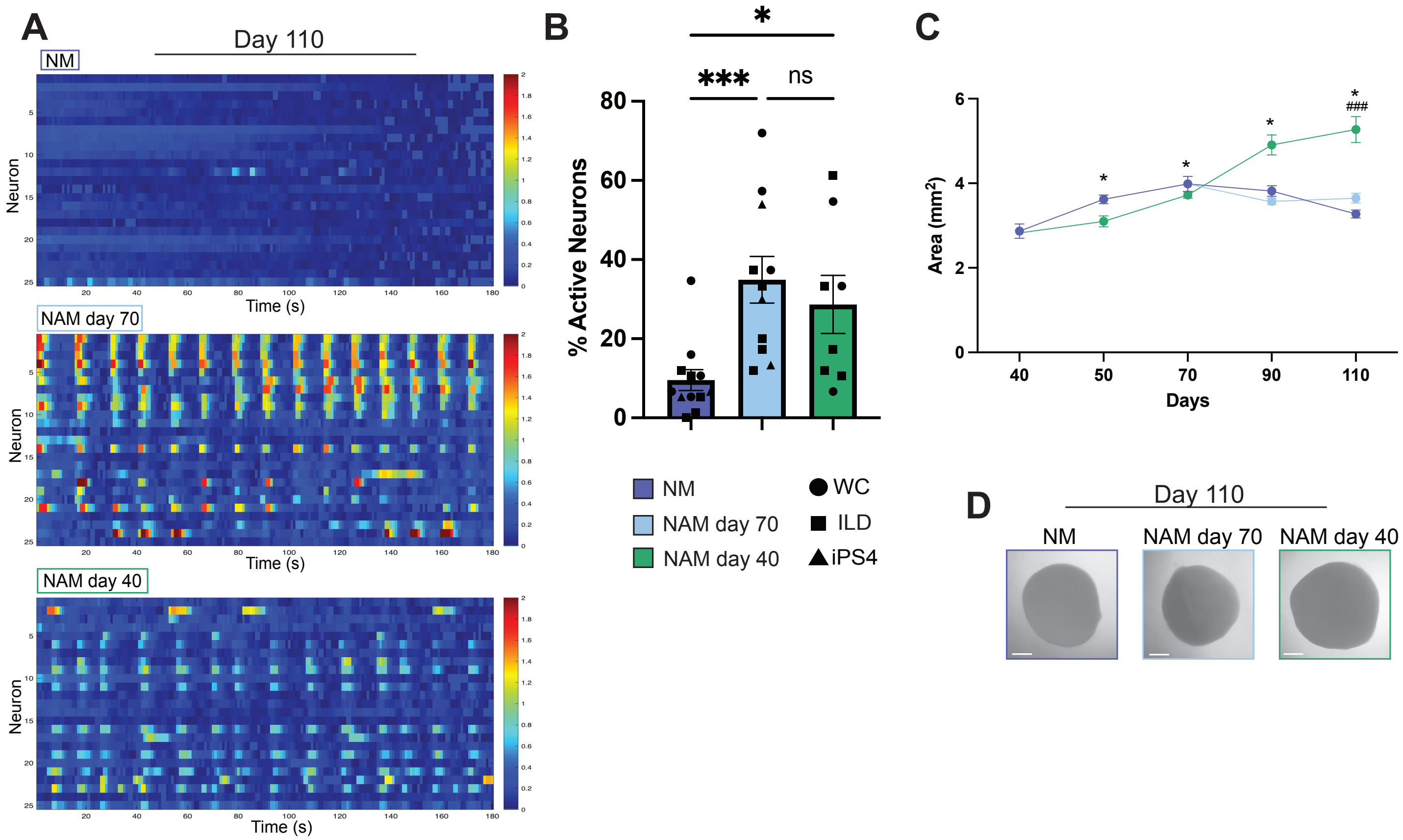


**Figure S4. Calcium imaging of COs.** (A) Representative heatmap with ΔF/F traces of NM-COs over 180s for each condition (NM, Neuronal Medium; NAM, Neuronal Activity Medium). (B) Percentage of active neurons in COs. Neurons are considered active if at least one calcium transient is detected in the 180s recording period. Each data point represents one CO, averaged across three separate fields of view (FOV). Data is from iPSC lines derived from three separate individuals. N = 8 – 12 COs per experimental condition, collected from 3 independent iPSC lines. **** $p \leq 0.0001$, Kruskal-Wallis test with Dunn's multiple comparison. Error bars represent mean ± SEM. (C) Longitudinal area measurements of COs from days 40 – 110. n = 22 – 38 COs per timepoint and per experimental condition, collected from 3 independent iPSC lines. * $p \leq 0.05$ between NAM day 40 and NM. ###, $p \leq 0.001$ between NAM day 70 and NAM day 40. Two-way ANOVA followed by Tukey's post-hoc multiple comparisons test. Error bars represent mean ± SEM. (D) Representative brightfield images of day 110 COs in three different experimental conditions. Scale bar is 50μm.

Figure S5

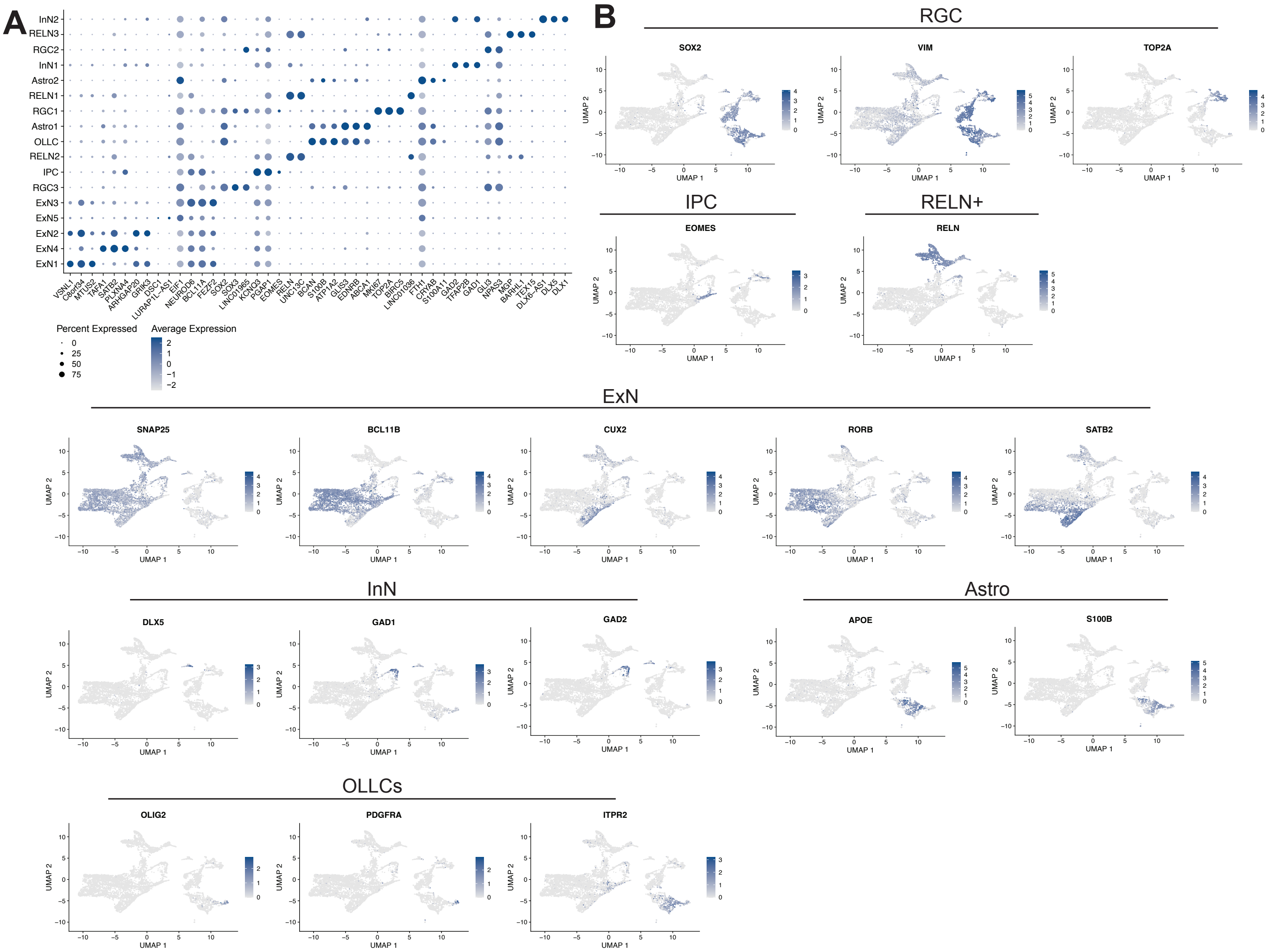


**Figure S5. scRNAseq cluster identification.** (A) Dot plot displaying top 3 differentially expressed genes per cluster. (B) Feature plots displaying expression of cluster-specific canonical marker genes.

Figure S6

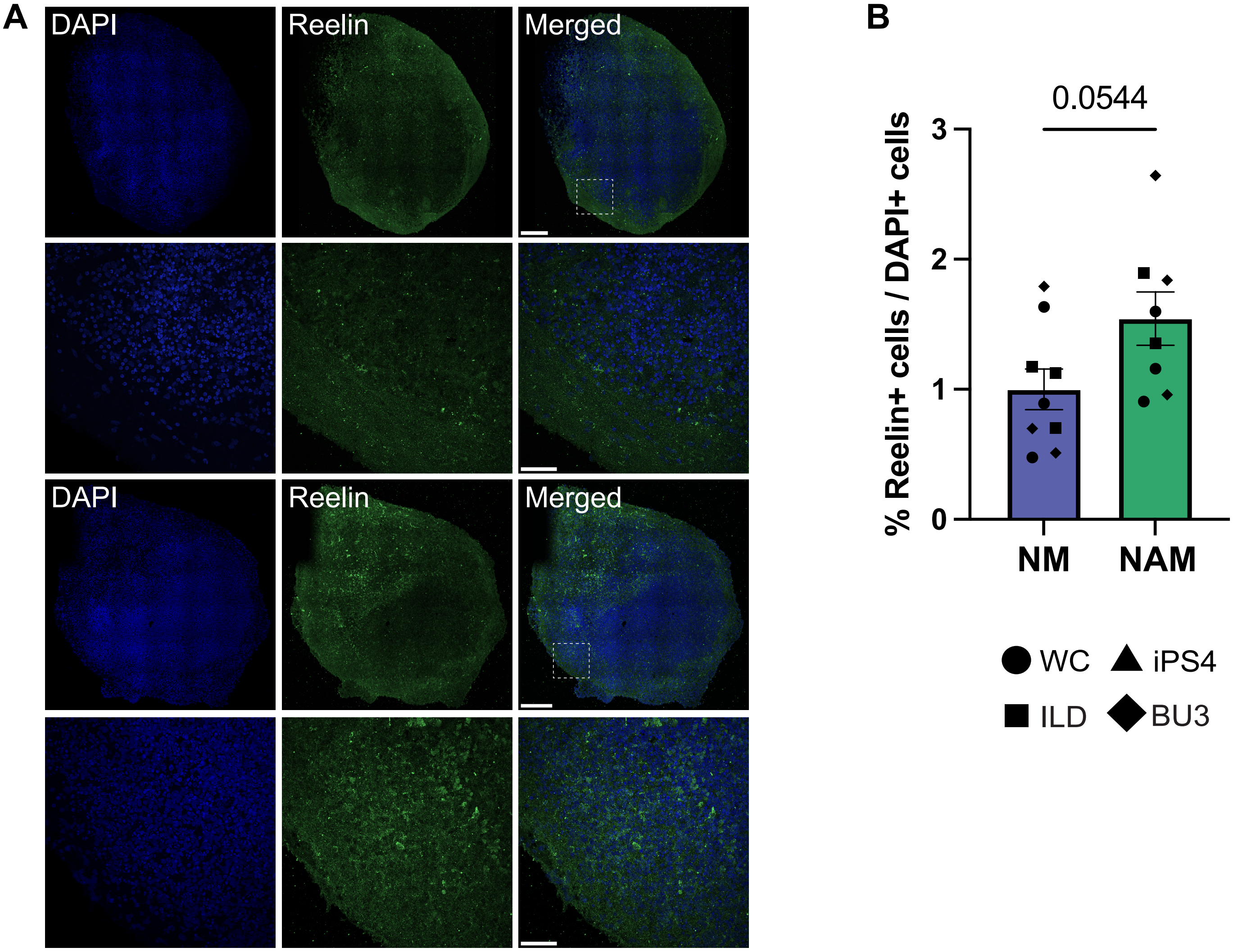


(A) Representative immunofluorescence images of day 110 OCOs stained with reelin. Scale bar 200μm, 50μm inset.
(B) Quantification of the proportion of Reelin+ cells, normalized to DAPI and quantified with QuPath v0.6.0. Each data point represents one OCO, averaged across three slices, and each shape represents a separate cell line. N = 12 OCOs. Error bars represent mean ± SEM
